# Association between theta-band resting-state functional connectivity and declarative memory abilities in children

**DOI:** 10.1101/2025.01.10.632362

**Authors:** Soléane Gander, Coralie Rouge, Anna Peiffer, Vincent Wens, Xavier De Tiège, Charline Urbain

**Affiliations:** Université Libre de Bruxelles (ULB), ULB Neurosciences Institute (UNI), Center for Research in Cognition & Neurosciences (CRCN), Neuropsychology and Functional Neuroimaging Research Unit (UR2NF), Brussels, Belgium; ULB, UNI, Laboratoire de Neuroanatomie et Neuroimagerie translationnelles (LN2T), Brussels, Belgium; ULB, Hôpital Universitaire de Bruxelles (H.U.B.), CUB Hôpital Erasme, Department of Translational Neuroimaging, Brussels, Belgium

**Keywords:** Declarative memory, magnetoencephalography, resting-state functional connectivity, theta oscillations, childhood

## Abstract

Declarative memory formation critically relies on the synchronization of brain oscillations in the theta (4–8 Hz) frequency band within specific brain networks. The development of this capacity is closely linked to the functional organization of these networks already at rest. However, the relationship between theta-band resting-state functional connectivity and declarative memory abilities remains unexplored in children. Here, using magnetoencephalography, we examined the association between declarative memory performance and pre-learning resting-state functional connectivity across frequency bands in 32 school-aged children. Declarative memory was assessed as the percentage of correct retrieval of 50 new associations between non-objects and magical functions, while resting-state functional connectivity was measured through power envelope correlation of the theta, alpha, low and high beta frequency bands. We found that stronger theta-band resting-state functional connectivity within occipito-temporo-frontal networks correlated with better declarative memory retrieval, while no correlation was observed in the alpha and beta frequency bands. These findings suggest that the functional brain architecture at rest, specifically involving theta-band oscillations, supports declarative memory in children. This mechanism may facilitate the subsequent rapid transformation of sensory input into visuo-semantic representations, highlighting the critical role of theta-band connectivity in early cognitive development.

## 1 Introduction

Children, particularly at school age, have to quickly learn large amounts of novel information about their environment (Hart et al., 2007). The declarative memory (DM) system is particularly at play to this aim, as it allows children to acquire and store broad conceptual representations and explicit knowledge about facts and events (Eichenbaum, 1997; Hart et al., 2007; Squire, 2004; Tulving, 2002).

Over the past decades, functional magnetic resonance imaging (fMRI) studies have characterized the brain regions underlying DM processes in adults (review in Kim, 2011). They demonstrated the key role of (para-)hippocampal and neocortical (e.g., prefrontal or temporo-parietal) interactions in the storage and the ‘binding’ of DM representations (Jenkins & Ranganath, 2010; Moscovitch et al., 2016; Ofen, 2012; Squire, 2004; Staresina & Davachi, 2010; Tang et al., 2018; Tulving, 2002; Vincent et al., 2006). In particular, it was suggested that hippocampo-neocortical functional connectivity (FC) processes allow the binding of *a priori* arbitrarily related elements (e.g., face-or object-name; object-function; words-meaning) into unique memory traces or *engrams* as well as their transfer into pre-existing memory systems (Cohen et al., 1997; Eichenbaum, 2001; Squire, 2004).

During childhood, these hippocampo-neocortical networks are continuously maturing (Blankenship et al., 2017; Ghetti & Bunge, 2012; Menon, 2013; Ofen, 2012; Uddin et al., 2011), in parallel with the development of complex learning and cognitive functions such as DM. Understanding how the functional brain architecture is related to such abilities in children aims at providing new information about the complex brain networks that are essential for learning and memory processes across development. Furthermore, it paves the way for a better understanding of the neuro-functional processes that may be altered in specific learning disabilities or neurodevelopmental disorders.

Interestingly, studies have shown that, already at rest (i.e., in the absence of explicit or goal-directed task practice), the functional organization of memory-related networks could play a critical role in the development of learning abilities (Gerraty et al., 2014; Wang, LaViolette, et al., 2010). Resting-state FC (rsFC) is considered as a marker of functional brain network integrity or efficiency (Smith et al., 2009) and has been shown to be predictive of functionally related performance or abilities in adults (Schlaffke et al., 2017; Stillman et al., 2013; Wang, Negreira, et al., 2010). Regarding DM, few fMRI studies reported an association between interindividual variability in episodic memory performance (i.e., DM for events) and rsFC across DM-related regions of interest (ROI) such as the hippocampus and cortical regions (e.g, cingulate, precuneus, retrosplenial cortex, inferior parietal lobule), in adults (Persson et al., 2018; Vincent et al., 2006; Wang, LaViolette, et al., 2010; Wang, Negreira, et al., 2010). Similarly, three fMRI studies showed that hippocampal ROI-based rsFC were correlated with DM abilities in pre-school-aged children (i.e. aged 4 to 6 years) and in adolescents (Geng et al., 2019; Riggins et al., 2016; Warren et al., 2021). Precisely, these studies showed that episodic memory performance is associated with hippocampal-dependent rsFC processes in networks involving the precuneus, the superior frontal gyrus (SFG), the superior temporal gyrus (STG) (Riggins et al., 2016) as well as the orbital frontal gyrus (OFG) (Geng et al., 2019) in young children and the inferior parietal lobule (IPL) in adolescents (Warren et al., 2021).

Still, none of the above-mentioned studies have investigated DM-related rsFC processes using a broader brain approach, which do not constrain their purview to the study of hippocampal-dependent rsFC. In addition, due to the sluggishness of fMRI responses (Heeger & Ress, 2002; Logothetis, 2008), these past studies could not investigate theta frequency band (4–8 Hz) oscillatory activity, which has been reported as critical for learning and memory processes (Fell & Axmacher, 2011; Solomon et al., 2017; Staresina & Wimber, 2019). Accordingly, using intracranial or magnetoencephalographic (MEG) recordings, previous studies reported the involvement of long-range theta-band coupling between medial temporal lobe (MTL) and prefrontal cortex (PFC) in successful DM encoding or retrieval in adults (Anderson et al., 2010; Backus et al., 2016; Kaplan et al., 2014) and children (Johnson et al., 2022). Theta-band oscillatory coupling between fronto-temporal brain regions was thus understood as the possible electrophysiological mechanism that enables the transfer of information during memory formation.

As MEG measures neuronal activity directly at the millisecond (ms) level and with a good spatial resolution, it has the ability to analyze long-range functional brain connectivity processes in specific frequency bands (Engel et al., 2013; Hipp et al., 2012). MEG thus provides an exquisite opportunity to investigate how long-range theta-band rsFC prior to DM learning relates to subsequent DM performance in school-aged children, as previously shown in the context of procedural learning and working memory tasks (Barnes et al., 2016; Van Dyck et al., 2021). Based on existing fMRI studies exploring the link between interindividual rsFC and DM performance in young children (Geng et al., 2019; Riggins et al., 2016) and the established role of theta-band oscillations in DM processes (Backus et al., 2016; Fell & Axmacher, 2011; Kaplan et al., 2014), we hereby hypothesized that stronger theta-band rsFC within temporo-frontal brain networks is associated with better DM performance at school age.

## 2 Methods

### 2.1 Participants

Thirty-seven typically developing children aged from 7 to 12 years old participated in this study. All children were right-handed, as measured by the Edinburgh Handedness Inventory scale (Oldfield, 1971) and native French speakers with self-reported normal or corrected vision. They had no prior history of neurological/psychiatric issues or learning, cognitive or language disabilities. The “Sleep Disturbances Scale for Children” (SDSC) (Bruni et al., 1996) allowed us to ensure that all children had normal sleep habits over the preceding month and the 3 nights preceding the experiment (“St Mary’s Hospital Sleep Questionnaire”) (Ellis et al., 1981). Vigilance was also measured before the MEG recording sessions (see below) using the psychomotor vigilance task (PVT, 5 minutes-duration version) (Dinges & Powell, 1985) to control for potential variations in sustained attention abilities between participants (Chun & Turk-Browne, 2007).

Overall, five children had to be excluded from the analyses due to excessive movements during MEG recordings (n = 3), poor sleep quality (n = 1) or poor vigilance (n = 1). The final sample therefore included thirty-two school-aged children (15 females and 17 males, mean age ± SD: 10.0 ± 1.1 years).

The study was approved by the Ethics Committee of the HUB-Hôpital Erasme (Brussels, Belgium; Reference: P2018/335). Participants and their parents gave written informed consent.

### 2.2 Material and design

#### 2.2.1 Material

Complete details regarding the learning material features and procedure can be found in Urbain et al. (2013, 2016).

Briefly, one hundred 2D colored drawings of unknown non-objects (NO) were used in the DM task (figure 1A). All NO were randomly split between 50 to-be-learned NO and 50 control (i.e., not learned) NO. Each to-be-learned NO was randomly associated with a magical (imaginary) function that had to be learned by the participants (e.g., “paints the sky in all colors”, “opens all doors”, “stops the rain”, etc.). All definitions were in French and included 3 to 7 words.

**Figure 1.**
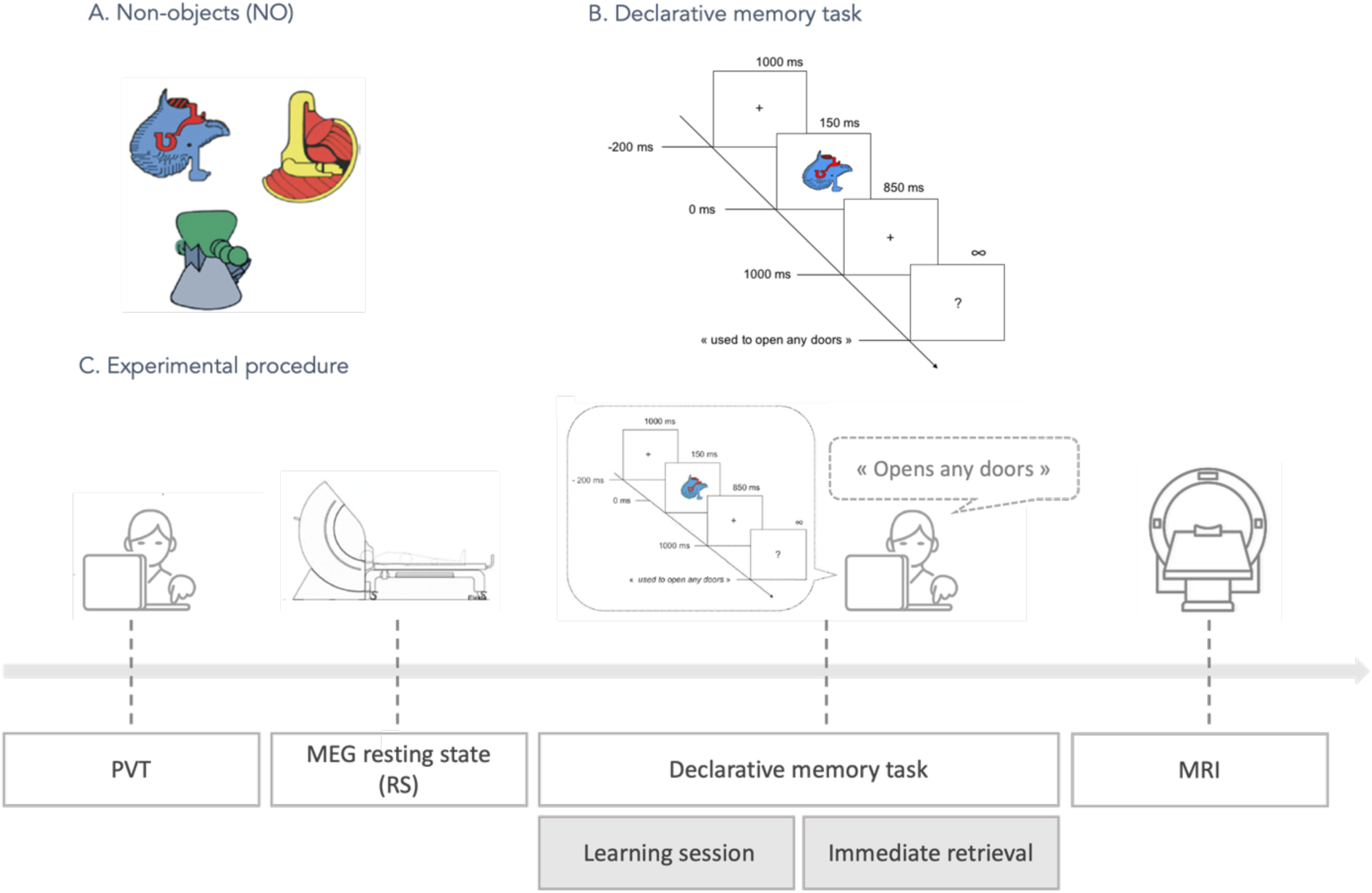
Experimental task and procedure design. **A.** Sample illustrations of the 50 non-objects (NO) used. **B.** Declarative memory task: At each trial, children were asked to provide the definition of the NO presented on the screen, or skip it if the NO was not previously defined. Responses had to be given after the appearance of a question mark (1 s after stimulus onset). **C.** Experimental procedure: After a psychomotor vigilance task (PVT) and two 5-min resting-state MEG recordings sessions, children undergo the declarative memory task separated into a learning session and an immediate retrieval session. A final structural high-resolution brain 3D T1-weighted magnetic resonance image (MRI) is acquired.

#### 2.2.2 Experimental design

The experimental protocol lasted for a half day and occurred as followed (see figure 1C for details). Participants first performed the PVT (5 minutes-duration version) (Dinges & Powell, 1985), during which they had to press a response button as fast as possible each time a digital counter stimulus was presented. The mean reciprocal reaction time (RRTs = 1/Reaction Time(s)) was used as vigilance score as recommended by Basner & Dinges (2011). Then, they underwent two successive 5 minutes resting-state (RS) MEG recordings, which aimed at characterizing pre-learning rsFC processes in each individual. To do so, children were asked to lay down and remain as still as possible in the MEG scanner while focusing on a fixation cross placed on the ceiling of the magnetically shielded room. Immediately after, DM performance was assessed in a behavioral session based on the DM task previously developed by Urbain et al. (2013), which lasted approximately for one hour and included a forty-minute learning session and a fifteen-minute immediate retrieval session. During the learning session, children had to learn the magical functions associated with 50 NO objects (i.e., to-be-learned NO) until they reached a learning criterion of 60% correctly learned associations. If the participant did not reach this criterion, the learning session was repeated including only the presentation of the unlearned NO. Once the participant reached the criterion, the immediate retrieval session started. All 50 to-be-learned NO and 50 control NO were presented twice in a random order and children had to retrieve the functions associated with each of the 50 to-be learned NO and to correctly skip the 50 control NO. Each NO was presented for 150ms followed by an 850ms blank screen with a fixation cross, and then a question mark prompting the child to verbally provide aloud the correct object’s functions or say "skip" if unknown, with the question mark remaining on the screen until a response was given. The next trial began after a 1000ms inter-stimulus interval corresponding to a blank screen with a fixation cross (figure 1B). A performance score was calculated for each participant as the percentage of to-be-learned-NO correctly recalled during the immediate retrieval session. The protocol ended by the acquisition of a structural 3D T1-weighted brain magnetic resonance image (MRI) to allow for MEG source reconstruction.

### 2.3 Data acquisition, preprocessing and analyses

#### 2.3.1 MEG and MRI data acquisition

MEG data (signals band-pass filtered at 0.1–330 Hz and sampled at 1 kHz) were recorded inside a magnetically shielded room (Maxshield, MEGIN, Helsinki, Finland; see De Tiège et al. (2008) for details) using a 306-channel whole-scalp neuromagnetometer (MEG) system (Triux, MEGIN, Helsinki, Finland) installed at the HUB–Hôpital Erasme. The head position of each participant was continuously monitored inside the MEG helmet using four head-tracking coils (Taulu et al., 2005). In addition, three landmark positions (left and right tragi and nasion) and at least 400 additional head-surface points (on scalp, nose, and face) were digitized using an electromagnetic tracker system (Fastrak, Polhemus, Colchester, VT, USA). MEG-compatible bipolar electrodes were used to monitor ocular, cardiac, and mouth muscle artefacts, placed vertically around the eyes (electrooculography, EOG), on the back (electrocardiography, ECG) and vertically on the chin (electromyography, EMG), respectively.

After the two 5-min resting-state MEG sessions, a structural 3D T1-weighted MRI scan was acquired in all participants (MRI, 1.5T, Intera, Philips, Best, The Netherlands) except six for which MRI acquisition could not be performed successfully. For each of these participants, we used a linear deformation of the structural MRI of an age-matched child to best match head-surface points, using the CPD toolbox (Myronenko & Xubo Song, 2010) embedded in FieldTrip (Donders Institute for Brain Cognition and Behaviour, Nijmegen, The Netherlands, RRID:SCR_004849) (Oostenveld et al., 2011).

#### 2.3.2 MEG data preprocessing

MEG data were filtered using the temporal extension of signal space separation (tSSS; correlation coefficient, 0.98; window length, 10s) (Taulu et al., 2005) to remove external environmental noise and correct for head movements (Maxfilter, MEGIN, Helsinki, Finland; version 2.2 with default parameters).

Independent component analysis (FastICA algorithm with dimension reduction to 30 and hyperbolic tangent non-linearity contrast) (Vigario et al., 2000) was applied to band-pass filtered (1–40 Hz) MEG signals to remove remaining ocular and cardiac artifacts. Components related to artefacts were visually detected and regressed out of the full rank data of each session (number of components removed: 3.5 ± 0.8, range: 2–5). The cleaned MEG data were then filtered into the theta frequency band (4–8Hz), which was chosen for its specific role in DM processes (Backus et al., 2016; Fell & Axmacher, 2011; Kaplan et al., 2014) and in the alpha (8-12Hz), low beta (12-21Hz) and high beta (21-30Hz) bands to control for the specificity of the theta-band results.

#### 2.3.3 MEG source reconstruction

Participants’ structural brain MRI was preprocessed to compute the MEG forward model, which is necessary to proceed with MEG source reconstruction. The brain MRI was anatomically segmented using the FreeSurfer software (Martinos Center for Biomedical Imaging, Massachussetts, USA) (Fischl, 2012). MEG functional and brain MRI structural data were coregistered manually using the digitized fiducials and head-surface points (Mrilab, MEGIN Helsinki, Finland). A volumetric source grid (cubic with 5 mm edges) was defined in the Montreal Neurological Institute (MNI) template MRI and deformed onto each subject’s MRI using a non-linear deformation in Statistical Parametric Mapping Software (SPM12, Wellcome Trust Centre for Neuroimaging, London, UK) (Ashburner & Friston, 1999). The MEG forward model was then computed using the one-layer boundary element method implemented in the MNE-C suite (Martinos Center for Biomedical Imaging, Massachusetts, USA) (Gramfort et al., 2014).

Neuronal source activity in each frequency band was reconstructed using minimum norm estimation (MNE) (Dale & Sereno, 1993). A 10-minute empty room recording preprocessed with SSS and filtered in each frequency band was used to estimate the noise covariance matrix. The MNE regularization parameter was determined using the consistency condition described in a previous study (Wens et al., 2015). The resulting three-dimensional dipole time series were further processed by projecting them onto their direction of maximum variance and obtaining their analytic signal via Hilbert transformation, as previously described by Sjøgård et al. (2019) and Wens et al. (2014).

#### 2.3.4 Functional connectivity analyses

Theta-band rsFC between each pair of sources was calculated through power envelope correlation (Brookes et al., 2011; Hipp et al., 2012; Wens et al., 2014) of these theta-band oscillatory source signals, which were orthogonalized beforehand to correct for spatial leakage (Brookes et al., 2012). This procedure was repeated for the other frequency bands (i.e., alpha, low beta and high beta bands). Power envelopes were also low-pass filtered at 1Hz before being correlated (Hipp et al., 2012). The temporal correlation between each pair of the envelope signals was calculated separately for each resting-state MEG recording (5-min) and then averaged over the two sessions to improve the stability of the power envelope correlation estimation, as recommended in previous studies (e.g. see Liuzzi et al. (2017)). This measure of connectivity was estimated at the connectome level using a customized structuro-functional parcellation of the human brain, including 75 nodes (MNI coordinates of all nodes can be found in memory atlas developed by Peiffer et al. (2021)). This parcellation involves 47 nodes from the Automated Anatomical Labeling (AAL) atlas (Tzourio-Mazoyer et al., 2002) and 28 additional nodes distributed over the whole brain that were selected from the literature for their partial or specific role in DM processes (Bastin et al., 2019; Ranganath & Ritchey, 2012; Takashima et al., 2009; Urbain et al., 2016). The resulting 75-by-75 connectivity matrices were calculated for each subject and for each frequency band yielding an image of their functional connectome in the frequency bands of interest (Figure 2A). To prevent potential asymmetries that could arise after pairwise orthogonalization, we followed the approach described in Hipp et al. (2012) and symmetrized the matrices by taking their average with their own transpose. To exclude a potential impact of power on functional connectivity estimates (Muthukumaraswamy & Singh, 2011), we calculated the variance of the source signal at each node with depth bias correction by noise standardization (Pascual-Marqui, 2002). The influence of signal power from each node pair on the power envelope correlation of the correction connection was suppressed alongside other covariates of no interest (see below) using multivariate regression modeling.

**Figure 2.**
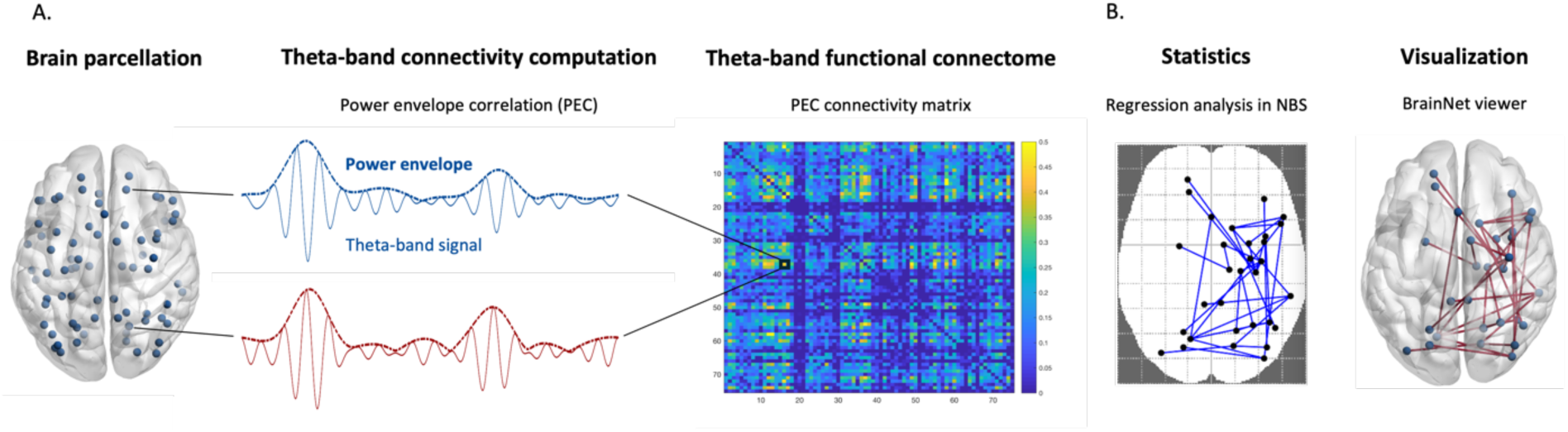
Functional connectivity and statistical analyses. **A.** Neuronal theta-band source signal was reconstructed for 75 nodes from a customized structuro-functional parcellation of the human brain. Theta-band rsFC between each pair of nodes was calculated through power envelope correlation (PEC) of the theta-band source signals. The resulting 75-by-75 connectivity matrices were calculated for each subject. **B.** The link between interindividual rsFC and behavioral retrieval performance was computed for each connection in the functional brain connectome, using a regression analysis implemented in network-based statistic (NBS). Significant brain networks were then visualized using the BrainNet Viewer Connectivity Toolbox.

#### 2.3.5 Statistics and reproducibility

Behavioral DM performance scores were calculated for each participant as the percentage of learned NO correctly recalled during the immediate retrieval session. We checked for outliers considered here as exceeding 3 median absolute deviations from the median DM performance score (Leys et al., 2013), removing them from the analyses where appropriate.

We used a regression analysis to infer statistical relationships between behavioral DM retrieval performance and MEG pre-learning theta-band rsFC for each connection across the connectome. To test the specificity of the theta frequency band, we also replicated these regression analyses with alpha-, low beta-and high beta-bands rsFC. Performance score was inserted as covariate of interest. Age, sex, and number of trials needed for each participant to reach the 60% learning criterion were used as covariates of no interest and were regressed out the analysis alongside node signal power (see above). The vigilance score (i.e., RRTs) was further added post-hoc as covariate of no interest in order to check for potential confounding contribution of vigilance performance to actual DM retrieval performance. A regression analysis with vigilance as the covariate of interest was also performed to confirm the findings of this post-hoc analysis. Positive regression coefficients indicated that higher rsFC was associated with better behavioral performance, while a negative regression coefficient indicated that higher rsFC was associated with poorer behavioral performance. The statistical significance of the regression coefficients was assessed using non-parametric network-based statistic (NBS) (Zalesky et al., 2010). NBS allowed us to identify network components, i.e., contiguous sets of brain connections with significant correlation between rsFC and behavioral performance, while keeping control of the Family-Wise Error Rate (FWER). A FWER-corrected *p*-value was assigned to each network component using permutation testing (n = 10000). To target strong correlations, a conservative univariate threshold of t ≥ 4.5 was applied. Significant brain networks (i.e., associated with *p*_corr_ < 0.05) were then visualized using the BrainNet Viewer Connectivity Toolbox (Xia et al., 2013) (Figure 2B).

## 3 Results

### 3.1 Behavioral performance related to declarative memory and vigilance

No participant was identified as a behavioral outlier based on DM performance score (i.e. percentage of correct responses for the learned NO). During the immediate retrieval session, children successfully recalled 64 ± 13% (mean ± SD percentage of correct responses) of the functions associated with the 50 learned NO. The number of trials needed to reach the criterion (60%) in the learning session was on average of 2 and did not exceed 3.

One child was excluded from the analyses due to excessively low vigilance (score exceeding 3 median absolute deviations from the median RRT score). The mean RRT over all children was of 2.70/s (SD = +-0.32/s).

No significant correlation was found between vigilance (RRTs) and DM performance (Pearson correlation test, *p* > 0.3), nor between age and DM performance or vigilance (Pearson correlation test, *p* > 0.3).

### 3.2 Link between theta-band resting-state functional brain connectivity and declarative memory performance

Regression analyses revealed significant positive correlations between pre-learning theta-band rsFC and subsequent DM performance that emerged within two, mainly right-lateralized, neural networks (NBS, *p*_corr_*s* < 0.02). The first network included 5 connections among 6 nodes, namely connections between the right superior occipital gyrus and the medial superior frontal gyrus, the right medial prefrontal cortex, the left orbital frontal gyrus and the right amygdala, and a connection between the medial superior frontal gyrus and the right inferior temporal gyrus (*p_corr_* = 0.001). The second network included a connection between the anterior part of the right inferior temporal gyrus and the right fusiform gyrus (*p_corr_* = 0.019; see all components regrouped in Figure 3, upper panel for the axial view and lower panel for the sagittal view). No negative correlation was found.

**Figure 3.**
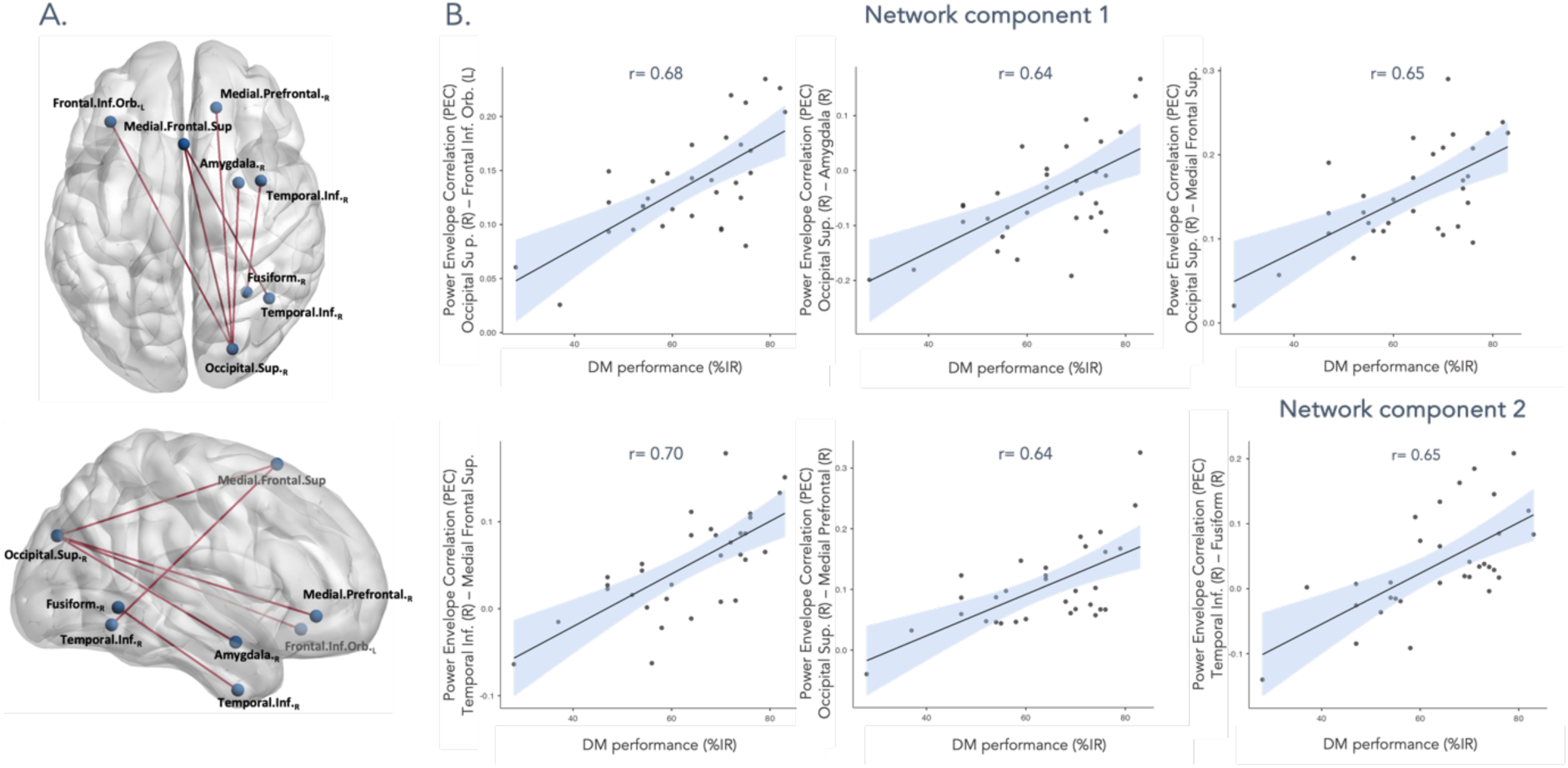
Significant correlations between pre-learning theta-band resting-state functional connectivity and subsequent declarative memory performance. **A.** Connections positively correlating with declarative memory (DM) performance, represented on the MNI brain (viewed from the top (up) or the right (down)) with nodes obtained from the customized structuro-functional parcellation. These plots were realized using the BrainNet viewer (Xia et al., 2013). **B.** Scatter plots for each significant connection from the two network components. L/R = left/right hemisphere, %IR= percentage of correctly recalled objects during immediate retrieval.

Repeating these analyses (post-hoc) with vigilance score as additional covariate of non-interest, we observed that the significant positive correlations between pre-learning theta-band rsFC and subsequent DM performance were reduced to two significant connections (*p*_corr_*s* = 0.021) including the right superior occipital gyrus connected to the left orbital inferior frontal gyrus, and the right inferior temporal gyrus connected to the medial superior frontal gyrus. Worth noticing, this later result was similar to the result obtained with our initial analysis when the NBS threshold was reduced (t < 4.5), suggesting that vigilance is not a primary contributor to the DM-related networks. This was further confirmed by the absence of significant correlation between vigilance score and theta-band rsFC (NBS *p*_corr_ > 0.05).

Complementary regression analyses in the alpha (8-12Hz), low-beta (12-21Hz) and high-beta (21-30Hz) frequency bands revealed no significant correlation between rsFC and DM performance (NBS *p*_corr_*s* > 0.05), suggesting that DM performance is specifically related to theta-band rsFC.

## 4 Discussion

This study aimed at better understanding how the functional architecture of the developing brain, sustained by theta-band cortical oscillations at rest, is associated with subsequent DM abilities at school-age.

Results showed that 7- to 12-year-old children with stronger theta-band rsFC within two mainly right-lateralized occipito-temporo-frontal networks were also better at retrieving newly learned DM information. More precisely, better DM performance was associated with stronger theta-band rsFC between (i) the right superior occipital gyrus, the medial and the left orbito-frontal cortices, the right amygdala and the right inferior temporal gyrus, but also between (ii) the right inferior temporal gyrus and the right fusiform gyrus. Altogether, these findings show that the strength of offline theta-band connections between specific brain network nodes influences subsequent DM performance in children.

While this study establishes a link between the functional organization of the resting brain of school-aged children and DM performance, similar associations have been observed with procedural memory at school-age (Van Dyck et al., 2021). Still, this past study revealed different underlying networks sustained by neural oscillations in other frequency bands. It was shown, using MEG, that procedural learning performance correlated with stronger pre-learning alpha-band rsFC within an interhemispheric sensorimotor network, encompassing the bilateral inferior parietal cortices and the primary somatosensory and motor cortices. In contrast, our study showed that DM performance was associated with stronger theta-band rsFC in two mainly right-lateralized occipito-temporo-frontal networks, suggesting a dissociation of underlying neural correlates between memory domains in the resting brain in school-age children. Supporting the hypothesis of a specific resting brain network for DM abilities, our results also indicate that the theta-band rsFC processes were not directly related to attentional abilities, as no significant correlation was observed between theta-band rsFC and the vigilance score.

Among other connections, better DM performance was associated with stronger rsFC between the right fusiform gyrus and the anterior part of the right inferior temporal gyrus in our study. As temporal areas are known to contribute to high-level processing and to the recognition of visual information (Geng et al., 2019), these results suggest that stronger functional connections between these regions at rest may have supported a better recognition of the learned NO during subsequent DM retrieval in our participants. This is in line with the known functional role of the right inferior temporal gyrus in the short-term storage and recognition processes of visual inputs, which allows to compare the incoming visual information with stored memory representations (Foxe et al., 2020; Ranganath et al., 2004) as well as of the right fusiform gyrus, which facilitates the transformation of the processed sensory input (e.g., visual input) into internal to-be-stored representations through the ventral stream of extra-striate visual systems (Kim, 2011). Moreover, increases in bilateral fusiform gyri activity have been frequently reported in the context of DM tasks in adults, particularly for successful encoding that helps later memory recognition processes (Garoff et al., 2005; Ofen, 2012). Hence, our results suggest that stronger rsFC within the right postero-anterior temporal areas may have improved the processing, as well as the encoding and/or the recognition of visual DM information in children.

This study also revealed that better DM performance was associated with stronger theta-band rsFC between bottom-up striate visual processing and top-down (pre)frontal brain areas. Amongst these top-down (pre)frontal regions, whose recruitment is known to depend on the content of the memory task (Wagner et al., 2001), rsFC involved the left orbito-frontal cortex (OFC) connected to the right superior occipital gyrus, the latter being mostly involved in the early stage of visual processing (Foxe et al., 2020; Geng et al., 2019). This finding aligns with previous studies reporting a role of the orbito-frontal regions in providing a rapid global estimate of the stimulus and its top-down control processing within relevant visual occipital regions (Bar, 2003; Bar et al., 2006; Engel et al., 2001; O’Shea & Walsh, 2006). Interestingly, this process may have been strengthened by the parallel functional connections of the right amygdala with the superior occipital gyrus observed in our results. Being particularly sensitive to rewarding or emotionally salient stimuli (Bar, 2003), we suggest that occipito-dependent brain processes related to the OFC and the amygdala support more efficient visual processing of learned information and, consequently, better interindividual DM performance. One could thus interpret that, from a phylogenetic perspective, these top-down mechanisms may have been progressively integrated into rsFC processes to enable the subsequent rapid transformation of salient sensory inputs into internal representations and their comparison with pre-existing memory traces.

The DM task used in our study not only required the recognition of a visual information (i.e., non-objects, NO) but also triggered the retrieval of associated episodico-semantic information (i.e. the NO’s magical functions) . This dual requirement is consistent with our results showing that a better DM performance was associated with stronger rsFC between occipital-temporal regions and medial (pre)frontal regions (i.e., the right mPFC and the superior medial frontal gyrus), which play a key role in the top-down access of semantic representations (Bokde et al., 2001; Jackson, 2021; Whatmough et al., 2002). Accordingly, our results suggest that stronger theta-band mPFC-dependent rsFC may have helped children in successfully retrieving the semantic (i.e., magical functions) information associated with the visual stimulus (i.e., NO). This interpretation is also supported by previous fMRI and MEG studies that have linked increased activity in mPFC and occipital-temporal regions to the optimal retrieval of semantic information, including information about an object’s function in children and adults (Urbain et al., 2013; Yee et al., 2010).

Worth noticing, despite the verbal semantic contribution of our DM task, which has often been described as mainly relying on the left hemisphere (Binder et al., 2009), the resting-state brain networks associated with DM performance in our study were predominantly right-lateralized. This lateralization is most likely due to the nature of our learning task requiring to bind idiosyncratic associations between novel visual, complex non-objects and their imaginary function and aligns with previous studies reporting rather right-than left-hemispheric processing in the context of complex picture learning tasks (Golby et al., 2001; Wagner et al., 1998), and especially in new situations for which no previous representation is available in long-term memory (Goldberg & Podell, 1994, 1995; Urbain et al., 2013).

Altogether, our results suggest that DM performance in children is supported by specific offline occipito-temporo-frontal theta-band functional brain connectivity processes, which may facilitate the encoding and the retrieval of new complex visuo-semantic representations (Staresina & Wimber, 2019). However, it is important to acknowledge certain limitations in this study. First, due to the correlational nature of our research design, we cannot establish causal relationships between rsFC and DM performance. Secondly, our study was based on a cross-sectional sample, limiting our ability to draw definitive conclusions about developmental trajectories and individual differences. Future studies using experimental designs, such as longitudinal investigations across multiple time points, would be valuable in assessing the stability and predictive value of theta-band rsFC for memory abilities.

Still, our results are in line with previous studies suggesting that theta-band oscillations specifically support the functional basis of long-range brain communication, in particular, between temporal areas and neocortical (mostly prefrontal) areas, required for memory formation in children (Johnson et al., 2022). Moreover, we did not observe a similar relationship between DM performance and rsFC in the alpha and beta frequency bands, strengthening the key specificity of theta-band FC processes for DM. Still, unexpectedly, MEG rsFC results did not include (para)hippocampal or medial temporal brain areas, despite their known role in learning and DM processes (Kim, 2011), including those involved in the specific task under used (see Urbain et al. (2016)). This is also surprising as theta-band oscillations have been repeatedly associated with hippocampo-dependent DM processes (Griffiths et al., 2021; Johnson et al., 2022; Staresina & Wimber, 2019) and hippocampal-related rsFC has been previously highlighted as a possible support for DM performance in young children (Geng et al., 2019; Riggins et al., 2016). Still, these latter studies were ROI hippocampal-based studies, which may have increased the chances of identifying an effect including medial temporal brain regions in past results. As a tentative interpretation, we suggest that rsFC processes are not key players for the specific process of binding memory traces, as this function is repeatedly associated with (para)hippocampal brain regions (Geng et al., 2019). Rather, our MEG results indicate that pre-learning rsFC would specifically support sensory input transformation processes, which are underpinned by larger brain networks. Precisely, our data suggest that stronger theta-band rsFC processes would facilitate the rapid processing and transformation of visuo-semantic information into internal representations, in the context of a DM task at school age. Consequently, stronger theta-band rsFC within these networks could lead to a more efficient communication between these brain areas during DM processes and thus to efficient visuo-semantic DM retrieval performance. This idea fits with the hypothesis that resting-state networks would ensure responsiveness to possible future tasks (Deco & Corbetta, 2011) by forming a critical pathway for communication when relevant. Furthermore, the identification of stronger theta-band rsFC as a correlate of better DM performance provides valuable insights into the potential underlying mechanisms contributing to learning difficulties. This could offer potential support for the early and easy (resting-state-dependent) identification of children at-risk for declarative learning impairments and possible insights regarding clinical routines and procedures.

## Data availability

The data that support the findings of this study are available on request from the corresponding author and after acceptance by institutional authorities (Hôpital Universitaire de Bruxelles and Université libre de Bruxelles). The data are not publicly available due to ethical restrictions.

## Acknowledgments and funding

SG was supported by the Fonds pour la formation à la recherche dans l’industrie et l’agriculture [FRIA, Fonds de la Recherche Scientifique (FRS-FNRS), Brussels, Belgium]. XDT is Clinical Researcher at the FRS-FNRS.

The MEG project at the Hôpital Universitaire de Bruxelles and Université libre de Bruxelles is financially supported by the Fonds Erasme (Convention « Les Voies du Savoir », Brussels, Belgium) and by research grants from the FRS-FNRS (FRS-FWO Excellence Of Science (EOS) MEMODYN: 30446199 ; CREDIT DE RECHERCHE: n° 29149840). The authors would also like to warmly thank all the children and their parents for their participation.

## Authors contribution

S.G. : behavioral, MEG and MRI data analyses; interpretation of the results, writing -original draft

C.R. : MEG and MRI data analyses, revising the manuscript

A.P. : investigation, revising the manuscript

V.W. : software, MEG and MRI data analyses, interpretation of the results, writing - original draft, funding acquisition

X.D.T. : conceptualization, interpretation of the results, writing - original draft, funding

C.U. : conceptualization, interpretation of the results, writing - original draft, funding

